# Cortical evoked activity is modulated by the sleep state in a ferret model of tinnitus. A case study

**DOI:** 10.1101/2024.05.12.593782

**Authors:** Linus Milinski, Fernando R. Nodal, Matthew K.J. Emmerson, Andrew J. King, Vladyslav V. Vyazovskiy, Victoria M. Bajo

## Abstract

Subjective tinnitus is a phantom auditory perception in the absence of an actual acoustic stimulus. It affects 15% of the global population and can be associated with disturbed sleep, poor mental health and quality of life. To date, there is no effective treatment for tinnitus. We used adult ferrets exposed to mild noise trauma as an animal model of tinnitus. We assessed the phantom percept using two operant paradigms sensitive to tinnitus, silent gap detection and silence detection, before and up to six months after the mild acoustic trauma. The integrity of the auditory brainstem was assessed over the same period using auditory brainstem response recordings.

Following noise overexposure, ferrets developed lasting, frequency–specific impairments in operant behaviour and evoked brainstem activity. To explore the interaction between sleep and tinnitus, in addition to tracking the behavioural markers of noise–induced tinnitus and hearing impairment after noise overexposure, we evaluated sleep–wake architecture and spontaneous and auditory–evoked EEG activity across vigilance states. Behavioural performance and auditory–evoked activity measurements after noise overexposure suggested distinct degrees of tinnitus and hearing impairment between individuals. Animals that developed signs of tinnitus consistently developed sleep impairments, suggesting a link between the emergence of noise–induced tinnitus and sleep disruption. However, neural markers of tinnitus were reduced during sleep, suggesting that sleep may transiently mitigate tinnitus.

These results reveal the importance of sleep–wake states in tinnitus and suggest that understanding the neurophysiological link between sleep and tinnitus may provide a new angle for research into the causes of phantom percepts and inform future treatments.

## Introduction

Sleep is a state of vigilance when neural activity is mainly generated endogenously while the brain is relatively disconnected from the external sensory environment [1–3]. During wakefulness, percepts arising from stimulus–independent neural activity, such as tinnitus or hallucinations, are sometimes a reflection of neuronal malfunction ([4]; work of Ambroise Paré reviewed in [5–7]). In contrast, during sleep, phantom percepts, such as imaginary sensorimotor experiences during dreams, are considered normal. Very little is known, however, about the relationship between the natural processes that take place during sleep and the pathological processes that give rise to persistent phantom percepts. The most prevalent phantom sensation is subjective tinnitus, or tinnitus for short, which affects an estimated 15% of the population [8,9]. Tinnitus is commonly reported as the constant perception of a hissing, ringing or buzzing sound without any identifiable acoustic source [8], and is often associated with depression, anxiety [10–12] and disturbed sleep [13–21]. To date, there is no available cure for tinnitus.

Tinnitus is associated with functional changes in widely distributed brain regions, including both auditory and non–auditory areas [22–28], many of which exhibit a dramatic modulation in their spatiotemporal activity across wakefulness and sleep [29–31]. We recently proposed that the spatial overlap between brain areas affected by tinnitus and those showing sleep–state dependent neural activity may lead to competition between pathological and physiological drives determining cortical network activity [32]. If tinnitus–related activity persists across vigilance states, it may result in a state of partial arousal during sleep similar to that observed in some forms of insomnia and parasomnias [33–35], where the emergence of global and local activation interferes with natural sleep–wake dynamics, potentially causing sleep impairments. Yet, the possibility remains that local and global changes in brain activity across vigilance states [30,36] may, in turn, interfere with tinnitus–related activity. In particular, high–intensity sleep with high levels of cortical slow wave activity, prompted by cellular and network–level drives for recovery sleep [37], such as after a period of extended wakefulness [30,38,39], could potentially mitigate tinnitus temporarily. This leads to the intriguing hypothesis that a dynamic modulation of the phantom sound sensation occurs across the sleep–wake cycle, depending on the relative weighting of circadian and homeostatic drives.

The aim of this study was to develop a new animal model for tinnitus, which will enable addressing this relationship and then systematically assess the interaction between sleep and tinnitus. The ferret is an attractive animal model that, due to its long lifespan, enables longitudinal studies to be carried out. Ferrets further allow for detailed assessment of operant behaviour, and are a valuable model for hearing research in particular, with a hearing frequency range more similar to the human range than in case of rodents. We tracked behavioural markers of noise–induced tinnitus and hearing impairment in a ferret model over a period of six months after noise overexposure and, in parallel, assessed the sleep–wake pattern, as well as spontaneous and auditory–evoked EEG activity across vigilance states. Behavioural performance and auditory–evoked activity suggested distinct differences in the degree of tinnitus and hearing impairment between individuals. Animals developing tinnitus also exhibited sleep impairments, suggesting a link between the emergence of noise–induced tinnitus and sleep disruption. Finally, neural markers of tinnitus were reduced during sleep, suggesting that sleep may transiently mitigate tinnitus. Overall, these results highlight the potential of measuring natural brain state dynamics to investigate tinnitus and uncover new avenues for future treatments.

## Material and Methods

### Animals

All procedures were carried out in accordance with the Animal (Scientific Procedures) Act 1986 (amended in 2012) and authorised by a UK Home Office Project Licence following approval by the Committee on Animal Care and Ethical Review of the University of Oxford. Seven adult female pigmented ferrets (*Mustela putorius furo*) were used in this study. Female ferrets are considered more suitable for neuroscientific studies than males because the brain size is similar across adult individuals and the thinner skull and temporalis muscles allow cranial implants to be attached more easily [40–42].

A subset of four animals were implanted for chronic recordings but only three were used for long term assessment due to an acute clinical condition in one animal unrelated to the implant. The animals were chronically implanted with cortical EEG and EMG electrodes to assess sleep–wake architecture and spatiotemporal patterns of brain activity before and after noise overexposure. The longitudinal design of this study spanning more than nine months allowed each animal to act as its own control by comparing the results before and after noise overexposure.

Animals were housed in small groups of at least three ferrets in standard laboratory enclosures or large pens housing up to ten animals. Food and water were available *ad libitum* except during periods of operant behavioural testing, where water access was mainly limited to rewards received during behavioural testing. Periods of water access regulation lasted for a maximum of five consecutive days before at least two days of *ad libitum* access to water. While under water regulation, animals received performance–dependent amounts of water during the behavioural task, topped up in the form of mashed food puree to an amount of 60 ml/kg body weight per day, to ensure that animals maintained a body weight ≥ 85% of their free–feeding weight.

Animals were housed under a 15/9h light–dark cycle from mid–March until mid– November (summer light cycle) and otherwise under an 8/16 h light–dark cycle (winter light cycle) to mimic natural seasonal changes in light exposure [43]. To suppress oestrus, ferrets were routinely injected with Delvosteron (MSD Animal health, Proligestone 100mg/ml) one month before the change to the summer light cycle. Behavioural experiments were conducted predominantly during the light phase, and chronic EEG recordings took place only during the summer light cycle.

During the periods of continuous EEG recordings (each lasting 2–3 days), animals were single housed in a custom–made enclosure (LxWxH 60×60×70cm) within a double– walled sound–attenuated chamber. Animals had *ad libitum* access to water and food. Identical bedding and nesting material to their home cages were provided, and the lighting and temperature conditions were the same as in the home enclosure. Before the first chronic recordings, ferrets were progressively habituated to the recording enclosure and to the tethering used for EEG recording. Recording enclosures were cleaned after each recording period.

Body weight, fur appearance and social interactions were monitored weekly over the course of the experiment to exclude any general change in animal behaviour due to the surgical or noise overexposure procedure or the recording paradigms.

### Electrode implantation

The aseptic surgical procedure largely followed the methodology previously described in [41,44]. Anaesthesia was induced with a single intramuscular injection of medetomidine hydrochloride (0.022 mg/kg BW; Domitor, Orion Pharma) and ketamine hydrochloride (5 mg/kg; Narketan10, Vetoquinol). Anaesthesia was maintained using Isofluorane (1-3%) (IsoFlo, Abbot Laboratories) with 100% oxygen as a carrier. Atropine sulphate (0.06 mg/kg, s.c.; Atrocare, Animalcare) was administered to minimise pulmonary secretions along with dexamethasone (0.5 mg/kg, s.c.; Dexadreson, Intervet) to prevent cerebral oedema. Doxapram hydrochloride (4 mg/kg, s.c.; Dopram–V Injection, Pfizer) was administered to minimise respiratory depression. Perioperative analgesia was provided with buprenorphine hydrochloride (0.01 mg/kg, s.c.; Vetergesic, Sogeval) and meloxicam (0.2 mg/kg, s.c.; Metacam, Boehringer Ingelheim). Prophylactic antibiotics to prevent infections were administered during the surgery (Augmentin: co–amyxoclav 0.02 mg/kg i.v. every 2 hours; Bechaam) and once daily for five days after surgery (Xynulox: Amoxicillin trihydrate/Co– amixoclav, 0.1 mg/kg i.m.; Zoetis). Depth of anaesthesia, respiratory rate, ECG, and end– tidal CO_2_ were monitored and maintained throughout the experiment. The animal’s temperature was monitored using a rectal probe and maintained at 38°C using a homeothermic electrical blanket (Harvard Apparatus) and a forced–air warming system (Bair Hugger, 3M Health Care).

The ferret was placed in a stereotaxic frame, the eyes were protected with a carbomer liquid eye gel (Viscotears, Alcon Laboratories), and the skull was exposed. Custom–made wired headmounts (Pinnacle Technology Inc. Lawrence) for EEG recordings were attached to bone–anchored stainless steel screws in contact with the *dura mater*. They acted as EEG electrodes, which were positioned unilaterally over the right frontal (AP 4 mm, ML 4 mm) and occipital (AP 7mm, ML 5mm) cortical areas and over the cerebellum reference electrode). Similar configurations of EEG electrodes have been used to provide recordings suitable for vigilance state scoring across different mammalian species such as ferrets, rats and mice [36,45,46]. Two tip–blunted stainless–steel wires were placed into the nuchal muscle for electromyography (EMG). Wires and screws were secured to the skull surface and protected by covering with bone cement (CMW1 Bone Cement, DePuy CMW, Lancashire, UK). EEG head mounts were protected with accessible plastic enclosures secured to the bone cement. The temporal muscle was temporarily detached at the dorsal part to provide access to the skull so that the EEG electrodes could be fixed to it. At the end of the surgery and to restore its function, the muscle was repositioned over the low profile most lateral part of the cranial pedestal using resorbable sutures and covered with the skin that was sutured independently around the most medial externalised part of the cranial implant. To expedite recovery from anaesthesia at the end of the procedure, animals received Antisedan (atipamezole hydrochloride, 0.06mg/kg, s.c., Vetoquinol). A minimum two–week postsurgical recovery period was allowed prior to further procedures.

### Noise overexposure

Noise (one octave narrowband noise centred at 8 kHz, 98 dB SPL at ear level) was presented for 120 minutes unilaterally via an earphone (Sennheiser CX300 II earphone) attached by a silicone tube to the entrance of the right ear canal while the left ear was fitted with an earplug and silicone impression material (Otoform, Dreve Otoplastik) to minimise its sound exposure. Closed–field calibrations of the sound–delivery system were performed using an 1/8th–in condenser microphone (Brüel and Kjær) attached to the silicone tube. The procedure was carried under general anaesthesia (assessed by immobility and absence of the pedal reflex), which was induced through intramuscular injection of medetomidine hydrochloride (0.022 mg/kg; Domitor, Orion Pharma) and ketamine hydrochloride (5 mg/kg; Narketan10, Vetoquinol). Depth of anaesthesia and the respiratory rate were monitored throughout the procedure. Anaesthesia was maintained by injection of half of the initial dose after 60 minutes or when the animal showed signs of arousal. Body temperature was maintained at 38°C using a homeothermic monitoring system (Harvard apparatus). To reverse the effect of Domitor and expedite recovery from anaesthesia, animals received Antisedan (atipamezole hydrochloride, 0.06mg/kg, s.c.). Noise overexposure (NOE) took place during the light phase, and animals were given at least 48h of rest before any other procedure.

### Auditory brainstem response measurements

Auditory brainstem responses (ABRs) were obtained under anaesthesia (medetomidine/ ketamine as described in ‘Noise overexposure’, see above) using sterile subcutaneous monopolar needle electrodes (0.35 x 12 mm, MN3512P150, Spes Medica). Body temperature was maintained at 38°C using a forced–air warming system (Bair Hugger, 3M Health Care). Stimuli were presented monaurally (left and right in subsequent recordings) via earphones (Sennheiser CX300 II) inserted into the ear canal and fixed in place with silicone impression material (Otoform, Dreve). Auditory stimuli were generated using an RP2.1 Enhanced Real–time processor (Tucker Davies Technologies, TDT) with a sampling frequency of 100 kHz connected to a TDT PA5 programmable attenuator. The earphones were calibrated using SigCalRP TDT calibration software to generate compensation filters ensuring stable levels across a frequency range from 250 to 30,000 Hz. Click stimuli (rarefaction click trains, rectangular voltage pulse, 100 µV, low–pass filtered) were presented at a rate of 17/sec for 700 repetitions per level (40, 50, 60, 70, 80, 90 dB SPL). One octave narrow–band noise stimuli (NBN, centred around 1, 4, 8 and 16 kHz) with a 5 ms duration were presented at a rate of 21/sec for 700 repetitions per level–frequency combination.

Signals were recorded from two active subcutaneous electrodes, placed close to the left and right auditory *bullae*, respectively, and referenced to an electrode placed at the vertex of the skull. A ground electrode was placed on the back of the animal. The signal was routed to a low impedance preamplifier (TDT RA16PA) and headstage (TDT RA4LI) and recorded by an RZ2 Bioamp Processor (25 kHz sampling rate) controlled by BioSigRP software (TDT).

### ABR signal analysis

ABR thresholds were determined manually by an experienced experimenter through visual assessment of ABR traces. This was conducted under blind conditions (enabled through a randomisation process used to access the data) with respect to animal, stimulus and stage of the experimental timeline (baseline, one week after noise overexposure (NOE) (Post1), and six months after NOE (Post2)). Thresholds were defined as the lowest stimulus level where an ABR wave was present if corresponding waves were also present at higher sound levels. If no ABR wave was present for any sound level, the threshold was defined to be at 90 dB SPL (the highest sound intensity used).

Data analysis was performed offline based on the average ABR signals (averaged over 700 individual ABR traces) for each stimulus type. As a readout for the magnitude of the entire ABR signal across all waves, the root mean square (RMS) of the signal was calculated by applying the MATLAB function *rms* on the signal in the predefined response window, 1.6 ms to 4 ms, to include only the ABR signal. To account for longer response latencies at low sound levels, the response window was shifted by 0.16 ms for each 10 dB decrement. A level–response plot was computed for each animal, assessment and stimulus and the area under the graph was calculated.

### Operant silent gap detection

Ferrets were trained by operant positive reinforcement using water as a reward to carry out a silent gap–detection task in an arena as described in previous work [47,48] (Fig S1A,B). The setup consisted of a circular arena (radius, 75 cm) housed in a double–walled sound– attenuated room. Animals were trained to initiate a trial by licking a spout, which activated infrared sensors on a platform at the centre of the arena. This ensured that the animal was facing the loudspeaker location at 0° azimuth at the time of sound delivery. Licking the central spout triggered the presentation of one of two types of sound stimuli through a single loudspeaker (Audax TW025MO). The two stimulus types were either a continuous sound or the same sound including four silent gaps. Following stimulus presentation, the animal had to leave the central platform and approach a peripheral response location at 30° to the left in ’gap trials’ and 30° to the right in ’no gap trials’ to obtain a water reward. Both types of stimuli were pseudo randomly balanced to avoid response bias to either location. There was no time limit for the animals to respond. Incorrect responses were not rewarded. After an incorrect response, trial initiation triggered the identical sound stimulus up to two more times (correction trials) before a new stimulus was presented. Correction trials were not included in the data analysis.

Sound stimuli were generated by TDT System III hardware. The paradigm was controlled by a custom MATLAB program that registered the position of the ferret at the arena -centre and response locations, presented the stimuli and delivered the rewards accordingly. Sound stimuli were broadband Gaussian noise bursts (BBN, 30 kHz lowpass) and one octave narrowband noise bursts (NBN) centred at 1, 4, 8, and 16 kHz. In gap trials, four equally spaced, identical silent gaps were introduced in the stimulus. Across trials, the length of these gaps varied from 3, 5, 10, 20, 50, 100 to 270 ms in duration. Stimuli were generated *de novo* for each trial, cosine ramped with a 10 ms rise/fall time and had a total duration of 2080 ms. All stimuli were filtered using the inverse transfer function of the loudspeaker to obtain stable sound intensity levels across the presented frequencies at 76±5 dB SPL.

Animals were tested twice daily in blocks of five consecutive days separated by at least two days of *ad libitum* access to water. Within each session, gap lengths were randomised across gap trials. Stimulus centre frequencies were identical across trials within a given session but varied between sessions to obtain approximately equal numbers of trials for all stimuli (1, 4, 8 and 16 kHz NBN and BBN). Procedural training, not included in the analysis, was provided using only the longest gap length (270 ms) until the animals reached ≥80% correct responses in two consecutive sessions, after which they were tested using all gap lengths.

Animals had to complete 600-1000 trials for each stimulus type. Analysis was based on the average performance for each session. Trials with response times of >5 seconds were excluded from further analysis. Sessions with few trials (more than 5 gap lengths each with <5 presentations) were also excluded from the analysis.

Since a constant phantom sound can fill a silent gap in a presented sound stimulus (Fig. S1A), tinnitus may create a bias towards detecting non-gap sounds. Consequently, lower FA and hit rates may occur with tinnitus, effectively compensating for each other when *d’* is calculated. Therefore, we used hit rate or the proportion of correct gap responses rather than calculating *d’* to quantify behavioural performance. Main effects on hit rate were estimated by fitting a GLMM (target distribution: normal, link: identity) on approximately normally-distributed hit rate data (repeated measures: gap length, testing session, stimulus frequency). Note that the number of gap and no gap trials was equal to prevent animals from developing a bias to one side in the paradigm. Therefore, no gap and gap trials contributed equally to statistical analysis, and the contribution of each gap trial was equal.

For analysis of silent gap detection thresholds, a sigmoid function (*R P (2022). sigm_fit (mathworks.com/matlabcentral/fileexchange/42641–sigm_fit, MATLAB Central File Exchange)* was fitted on hit rate (between 0 and 1) vs gap length, calculated for each animal, stimulus (1, 4, 8 and 16 kHz NBN and BBN) and condition (baseline, Post1 (1 week post NOE), Post2 (6 months post NOE)). Thresholds were defined as the gap length in closest proximity to a hit rate of 0.5 (chance level) on the fitted function. Fits with slopes at threshold of more than 10 times the median slope across all samples were excluded from the analysis (this applied to 2 out of 78 samples).

### Operant silence detection

Operant silence detection (modified from [49]) took place in the same testing arena as the silent gap detection paradigm, and trials were initiated in the same way. In the silence detection task, however, trial initiation triggered a light emitting diode facing the central platform (signalling the start of the trial) and one of three sound stimulus types: narrowband noise (NBN, one octave bandwidth with centre frequencies at 1, 4, 8, and 16 kHz randomised across trials), a sinusoidally–amplitude modulated BBN (AM stimulus, 100% modulation depth, 5 Hz modulation frequency) or silence (no sound). The proportions of trials in which these stimuli were presented were 50% for NBN, 30% for AM, and 20% for silence to ensure equal probability of reward in accordance with the following criteria. In AM and silence trials, responses to a sensor located at +30° relative to the central platform were rewarded with water, whereas in NBN trials responses to the –30° sensor were rewarded. There was no time limit for the animals to respond and a trial was completed whenever the animal responded to either of the two sensors.

During the training phase for this paradigm, reward probability for correct responses was gradually reduced from 1 to 0.7. After animals reached the performance criterion (>80% correct in two consecutive sessions), the paradigm was altered in that silence trials were never rewarded (’testing phase’ of the paradigm) to measure the animals’ performance without a further training effect. To keep the overall reward probability consistent with the training phase, the reward probability for non–silence trials was 0.9.

The animals completed a total of 1000 trials in this paradigm for each of the testing blocks, baseline and the two assessment blocks at different intervals after noise overexposure (Post1 and Post2). To reacquaint the animals with the paradigm before each testing block, they undertook the training paradigm again until reaching the performance criterion. While this retraining might have allowed animals to adjust to a new ’perceptual baseline’ after noise overexposure and therefore mask subtle effects of tinnitus or hearing impairment on performance, it ensured that any variations in performance over time were unlikely to be due to, e.g., the animal forgetting aspects of the paradigm, rather than an effect of NOE. To assess performance, hit rates were compared across stimuli and conditions (Baseline, Post1 and Post2). Sessions in which an animal completed less than 5 trials for one or more stimulus types were excluded from further analysis. Trials with long response times (>20s) were also excluded.

### Definition of indices for tinnitus and for changes in auditory brainstem responses

As a summary index for behavioural evidence for tinnitus a behavioural tinnitus index (TI, see Eq(1) below) was defined. The TI was calculated as the sum of three metrics obtained as indicators of tinnitus in the two operant tasks, by comparing the values in the baseline condition *(BL)* and Post NOE *(Post)*. Two of the metrics were based on the operant gap detection task (*M(cont)* and *M(thresh)*) and one on the silent detection task (*M*(silence).

For each metric, M(silence), M(cont), and M(thresh), positive values describe changes in line with tinnitus development (an impairment in silence detection, an increase in continuous sound detection and an increase in gap detection threshold, respectively). The behavioural tinnitus index (TI, see Eq(1)) is the sum of these metrics (Eq (2–4)), enabling the level of tinnitus and hearing loss experienced by each animal to be parametrised after NOE.

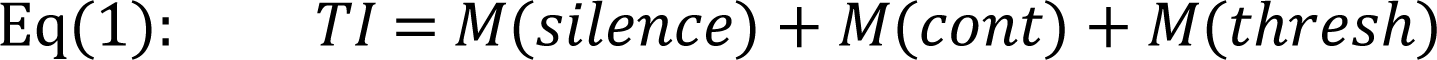

The metric for operant gap detection performance refers to the change in ability to detect the continuous sound (no gap) across all tested stimulus frequencies (Eq(2)) and to the change in gap–detection threshold across all stimuli (Eq(3)).

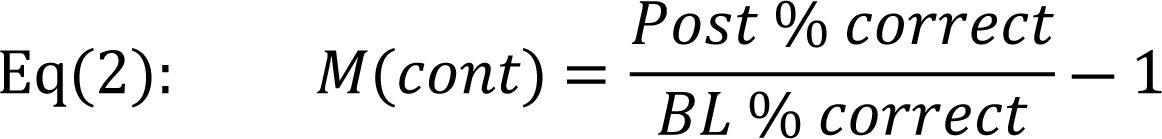

The metric for a change in gap detection threshold (Eq(2)) was based on a normalised performance value instead of thresholds corresponding to different gap lengths. To establish a normalised threshold, each gap length (3, 5, 10, 20, 50, 100, 270ms) was assigned a performance value based on the assumption that a threshold at 3ms corresponds to 100% performance and the remaining gap lengths correspond to evenly spaced decrements in threshold (85.71%, 71.43%, 57.14%, 42.86%, 14.29%). This approach for defining threshold changes results in a metric reflecting the direction of threshold change and its magnitude on a scale between 0 and 1).

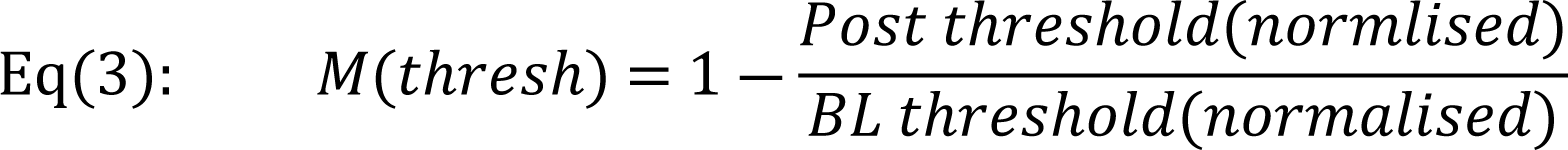

The metric for operant silence detection Eq(4) refers to the change in silence detection ability (percent correct in silence trials) after NOE.

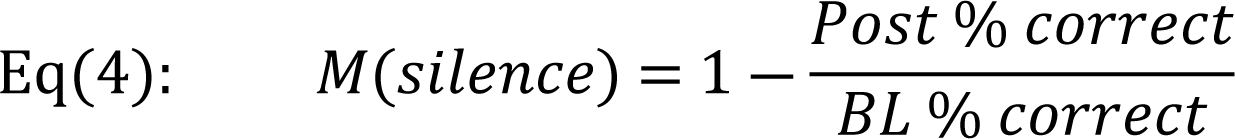

Thus, positive values describe the magnitude of a decrease in silence detection ability (and therefore evidence for tinnitus) and negative values describe the magnitude of an increase in silence detection ability. For example, a value of M (silence) = 0.2 would indicate a decrease in performance by 20% relative to baseline performance. The magnitude is between 0 and 1, the same scale as in the other defined metrics of the tinnitus index (Eq(2-3)).

### Index for changes in ABRs

Changes in ABRs were summarised using two metrics: first, changes in ABR thresholds relative to baseline (BL) and second, changes in ABR total magnitude relative to BL.

The metric for thresholds refers to the average change in thresholds across all tested stimuli (in dB), defined as the difference between BL and Post NOE values:

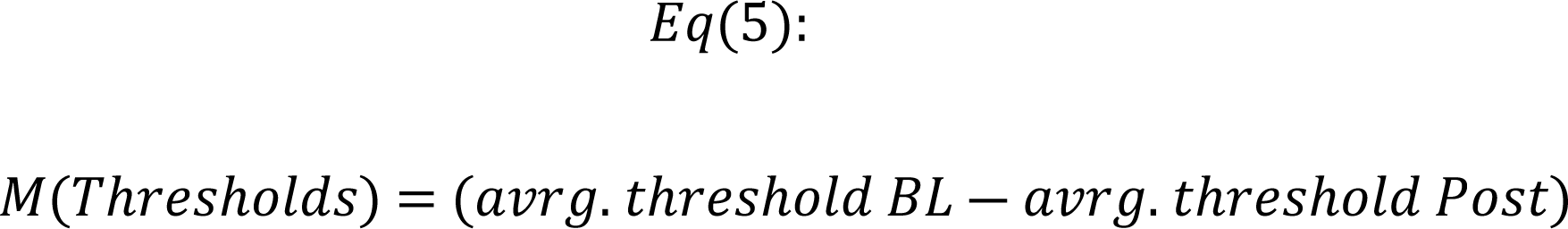

The metric for the ABR magnitude is defined as follows: first, the magnitude of the entire ABR was defined as the root mean square of the signal in a predefined response window (1.6-4ms, with a shift of 0.16ms for every decreasing step in sound level) following stimulus presentation. The total magnitude was then calculated for each animal and each stimulus by measuring the area under the level–response graph. The input ‘ABRmag’ used for the metric below is the average over total ABR magnitudes for all stimuli (1, 2, 4, 8, 16 kHz NBN and BBN) per animal. The metric for each animal is defined as

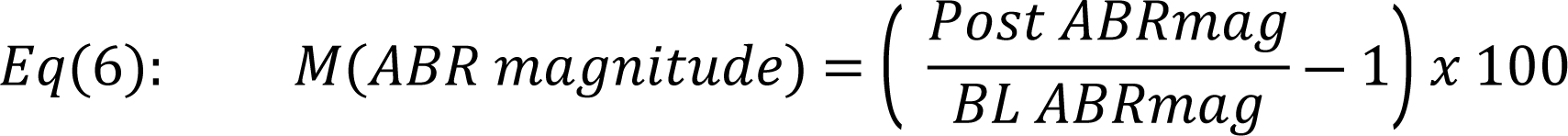

Positive values indicate the magnitude of a response increase (in %) whereas negative values indicate a reduced response (evidence for hearing loss).

### EEG signal processing and vigilance state scoring

Data acquisition was performed using a multichannel neurophysiology recording system (TDT). Cortical EEG was recorded from frontal and occipital derivations. EEG/EMG data were filtered between 0.1 and 100 Hz, amplified (PZ2 preamplifier, TDT) and stored on a local computer at a sampling rate of 1017 Hz, and subsequently resampled offline at 256 Hz. Signal conversion was performed using custom–written MATLAB (The MathWorks Inc.) scripts. Signals were then transformed into European Data Format (EDF). Vigilance state scoring was performed manually offline prior to spectral analysis and assessment of sleep architecture.

Manual vigilance state scoring was based on visual inspection of consecutive 4-s epochs of filtered EEG and EMG signals (SleepSign, Kissei Comtec Co). Frontal and occipital EEG and neck muscle EMG channels were displayed simultaneously to aid manual scoring and video recordings of the animal were consulted for further validation. Vigilance states were classified as waking (low–voltage desynchronised EEG with high level EMG activity), NREM sleep (presence of EEG slow waves, characterised by high amplitude and low frequency EEG, 0.5–4 Hz), REM sleep (low–voltage, mid–frequency EEG, 4.5-8 Hz, with a low level of EMG activity) or REM2 (low–voltage, high–frequency EEG, 8.5-20 Hz, with a low level of EMG activity. Epochs containing an EEG signal contaminated by artefacts (such as due to gross movements of the animal, eating or drinking) were excluded from subsequent analysis.

For each 24 h recording period, EEG power spectra were computed by a fast Fourier transform (FFT) routine for 4-s epochs (Hanning window), with a 0.25-Hz resolution (SleepSign Kissei Comtec Co). Artefacts in specific frequency bins that had remained unnoticed during manual data scoring were excluded offline. For each 0.25 Hz frequency bin of a given vigilance state during a 24 h recording period, the mean and standard deviation across all epochs was calculated based on a 500-iteration bootstrap. Values outside the mean ± 600 standard deviations across all epochs were excluded from the analysis (to exclude only extreme outliers). Exclusion in this case applied just to the specific frequency bin identified as an outlier. The objective of this was to prevent extreme outliers from distorting the spectral power estimate in particular frequency bins.

### Sound presentation during sleep and wakefulness

Animals were housed individually in a custom–made recording chamber on a 15/9 h light– dark cycle (summer cycle) as described above for the continuous EEG recordings. Frontal and occipital EEG and neck–muscle EMG recordings were obtained over approximately 48-72 hours (2-3 days) per recording session.

In the first 24 h per recording session, the animal was left undisturbed. Over the course of the second 24 h of the session, auditory stimuli were presented via a free–field loudspeaker installed on the ceiling of the double–walled sound–attenuated recording chamber above the custom–made enclosure. Sounds were one octave narrowband stimuli with centre frequencies of 1, 4, 8 and 16 kHz. Stimuli had a duration of 820 ms and included a silent gap of 38 ms in the middle of the stimulus. Stimuli were presented at 40, 50, 60 and 65 dB SPL (as measured at floor level at the centre of the enclosure). Inter–stimulus intervals had a random duration ranging from 10 to 42 s and each stimulus–level combination was presented 200 times. Stimuli were generated via MATLAB and produced using an RP2.1 Enhanced Real–Time Processor (TDT) and an Alesis RA150 Amplifier.

Stimulus presentation was controlled via a custom MATLAB script.

### Analysis of auditory evoked responses

Raw EEG data were transformed into a MATLAB compatible format (.mat) using the *tdtbin2mat* MATLAB function (provided by TDT). Evoked responses were analysed within a time window set between -0.5 s and +5 s relative to sound stimulus onset. Each stimulus presentation is referred to hereafter as a trial. Trials that fell into an epoch that contained an artefact (as defined during the manual vigilance state scoring procedure) were excluded from further analysis. To aid computing efficiency, the signal was downsampled by a factor of 2 (from an original sampling rate of 24414 Hz). Trials were then sorted into groups based on condition, stimulus frequency, level and vigilance state.

Due to marked inter-trial-variability in the EEG signal, conventional averaging across trials did not allow to define peaks and troughs of auditory evoked potentials (AEPs) reliably. To reduce the impact of noise on AEP detection, 20 bootstrapped means were drawn to serve as a representative signal sample for subsequent analysis, allowing for detection of peaks and troughs with minimal effect of noise while still reflecting the variability in the original dataset. Bootstrapped means were drawn from each group of trials (all trials for the same condition, vigilance state, stimulus and stimulus intensity), respectively. Peaks and troughs of evoked potentials were detected after smoothing each of those signals using a moving average of ∼8 points (implemented by the MATLAB *smooth* function). Peaks and troughs were automatically detected in a time window after stimulus onset that was predefined using a custom written MATLAB script and the *findpeaks* function. Time windows for early (R1), mid (R2) and late (R3) response components were defined based on the average latency of peaks and troughs in the respective animal, as the shape of the evoked potential was not uniform across animals. R1, R2 and R3 could each be identified in Ferret 2, R1 and R2 in Ferret 3, and just one response component, R1, in Ferret 1. Note that the response windows for R1, R2 and R3 components of the response were defined for each animal individually, depending on the latency of the respective deflection relative to stimulus onset. Response magnitudes for each response component were then defined as the difference between the maximum and subsequent minimum of the signal in the response window for the respective response component. For each animal, magnitudes of all response components were included (pooled, means±SEM) in the analysis.

Figures and illustration were produced by using MATLAB and MS PowerPoint.

## Results

### Ferrets develop long–term behavioural impairments indicative of tinnitus after noise overexposure

To establish an animal model of induced, persistent tinnitus, adult female ferrets (n=7) were tested in two operant paradigms sensitive to tinnitus (silent gap detection and silence detection) before (baseline, BL), starting one week (Post1) and, in a subgroup of ferrets (n=3) six months (Post2) after unilateral noise exposure (Fig 1A). The protocol for noise overexposure (2 hours of one octave narrowband noise centred at 8kHz at 98 dB SPL) was similar to protocols used for tinnitus induction in other animal models [50–52]. Noise overexposure triggers chronic tinnitus in human and animal models whereas salicylate models evoke reversible tinnitus [7,53,54]. Our experimental design was combined with regular assessment of auditory brainstem responses (ABRs, see Methods).

**Fig 1.**
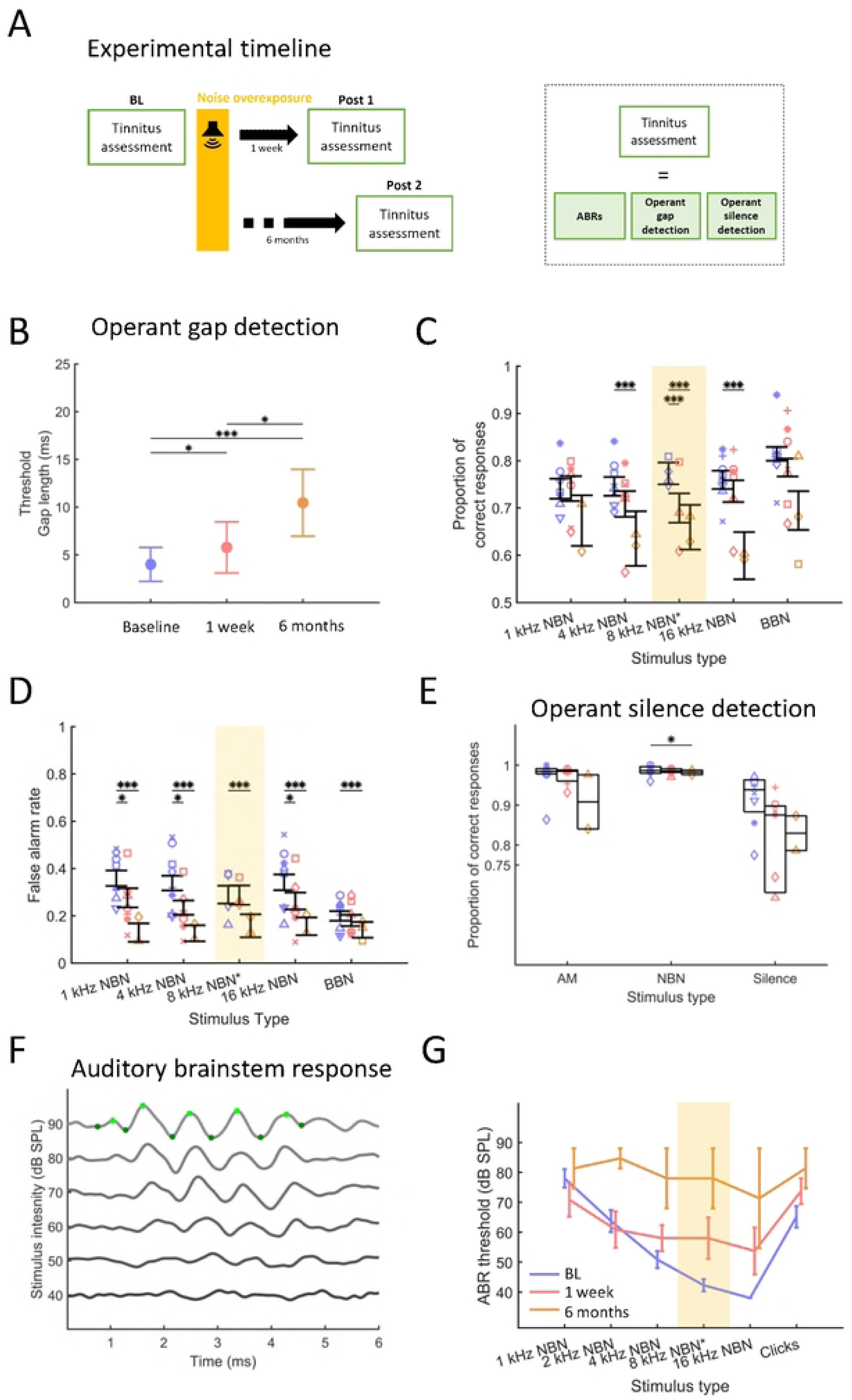
Behaviour and auditory brainstem responses are impaired after noise overexposure. **(A)** Experimental timeline. Animals were assessed in behavioural paradigms (operant silent gap detection, silence detection), and auditory brainstem responses (ABRs) were obtained. Assessments of all these metrics took place once under baseline conditions and on two occasions after noise overexposure (NOE), starting one week after NOE and within two months (Post1) and six months following NOE (Post 2). ABRs were always tested on one day in the first week following NOE and again 6 months later, whereas it took two months to complete behavioural testing at each of the three time points. (**B**) Mean operant gap detection thresholds across all narrowband (NBN) and broadband noises (BBN). Error bars represent standard deviation (SD) (**C**) Operant gap detection, proportion of gap trial correct responses for each one-octave NBN centred at 1, 4, 8, 16 kHz frequencies and BBN. Error bars represent 95% confidence intervals. (**D**) Operant gap detection, false alarm (FA) rate by stimulus type across time. FA = 1 - proportion of correct responses in no gap trials. Individual data points depict animal means. (**E**) Operant silence detection. Proportion of correct responses for AM, NBN and silence trials (see Fig S1D for setup). Box plots depict interquartile ranges across sessions. (**F**) Auditory brainstem response (ABR). Example traces showing 40-90 dB SPL click responses in one animal. Green markers depict peaks and troughs of ABR waves I to V, note the latency increase with intensity reduction. (**G**) ABR thresholds, defined as the lowest intensity where a significant response – local peaks of waves I-IV – were identified as in **F** by a trained experimenter under blind conditions. Data points in panels C to E represent the mean values from individual animals. Asterisks represent statistical significance, p values *<0.05, **<0.01, ***<0.001. Triangle, square and circle symbols in panels C, D and E represent the three cases in which cortical recordings were obtained during sleep (i,e. animals 1-3), also in Figs S1C and S2.

The primary operant paradigm was operant silent gap detection, which assessed the animals’ ability to detect short silent gaps in auditory stimuli [47] (Fig S1 A and S1B). Four animals were implanted for chronic recordings but only three were used for long term assessment. No behavioural differences were found between implanted (IM) and non-implanted (NIM) animals in baseline assessments and therefore behavioural data from the seven cases were analysed together (operant gap detection: BL threshold (IM) = 3.36±0.81 ms; BL threshold (NIM)=4.3±2.0 ms. Effect of group F_(1,32)_=0.04, p=0.85 (n.s.)).

In line with previous work on silent gap detection in ferrets [47], the animals’ performance declined as gap length was reduced (effect of gap length, GLMM, F_(7,6752)_=4489.37, p<0.001), an effect visible throughout the experiment (Figs 1A and S1C). After noise overexposure (NOE), silent gap detection ability was impaired, as indicated by the progressively lower percentage correct scores achieved at most gap lengths (Fig S2A) and by progressively increasing silent gap detection thresholds measured across all stimulus types (silent gap length at threshold, Baseline, BL: 4.0±1.78ms; Post1: 5.77±2.68ms; Post2: 10.45±3.5ms, means±SD; GLMM, BL vs Post1, β=1.69,t_(75)_=2.41, p=0.018; BL vs Post2, β=5.95,t_(75)_=11.68, p<0.001; Post1 vs Post2 β =5.46, t_(75)_=2.24, p=0.029. Figs 1B and S2A).

The initial impairment observed (GLMM, effect of condition, F_(2,941)_=32.31, p<0.001) was mostly due to reduced silent gap detection ability for 8 kHz narrowband noise burst (NBN) stimuli (probability of correct response, BL vs Post 1, 0.77±0.26 vs 0.7±0.27, β=-0.1, t_(943)_=-8.03, p<0.001), the NOE sound, whereas performance remained stable for other stimuli (p>0.1) (Fig 1C). Six months after NOE, the impairment was also present at neighbouring frequencies (4 and 16 kHz NBN) (4 kHz BL vs Post2, 0.75±0.27 vs 0.64±0.35, β=-0.04, t_(1319)_=-3.35, p<0.001; 16 kHz BL vs Post2, 0.76±0.26 vs 0.6±0.31, β=-0.14,t_(1359)_=-11.79, p<0.001. Fig 1C). The stimulus that differed most from the NOE sound in terms of frequency composition (1 kHz NBN) was the least affected across the course of the experiment (p>0.1). Accordingly, the stimulus that comprises a broad range of frequencies (BBN) was also less affected.

While silent gap detection ability progressively worsened after NOE, detection ability for no gap stimuli improved over time (Figs 1D and S2B). Specifically, ferrets showed statistically significant lower false alarm (FA) rates starting 1 week following NOE (Post 1) for 1, 4 and 16 kHz NBN (1 kHz, BL vs Post1, 0.36±0.15 vs 0.27±0.15, β=-0.1,t_(162)=-_2.66, p=0.01; 4 kHz, BL vs Post1, 0.34±0.15 vs 0.24±0.12, β=-0.12,t_(162)_=-2.84 p=0.01; 16 kHz, BL vs Post1, 0.34±0.16 vs 0.26±0.14, β=-0.08,t_(169)_=-8.75, p<0.001) stimuli, but not for 8 kHz NBN, the same stimulus as the NOE sound, and for BBN (p>0.1) (Fig 1D). This suggests that animals developed a temporary impairment in detecting both gap and no gap sounds consisting of 8 kHz NBN, whereas in frequency ranges adjacent to the NOE stimulus, they were more likely to respond to stimuli as if they were no gap sounds. In the longer term, six months after NOE, the FA rates were significantly lower for all stimuli, including 8 kHz NBN (p<0.001, Fig 1D) whereas the proportion of correct responses for gap stimuli was further impaired (Fig 1C). This improvement in FA rate over time, together with the gap detection impairment, suggest a tendency of the animals towards interpreting sounds with silent gaps as continuous sounds across all tested frequencies and could be an indication for tinnitus development.

To assess whether the impairment in silent gap detection ability after NOE could be attributed to diminished temporal resolution in auditory processing, the ferrets were tested in a silence detection paradigm [49]. They were tested in the same operant arena as for silent gap detection but had to discriminate NBN bursts of varying frequency compositions from ‘silence’, i.e. trials without any presented auditory stimulus, and from amplitude modulated (AM) BBN (see Methods & Fig S1D). As with operant silent gap detection, no behavioural differences were found between IM and NIM animals on silence detection and therefore behavioural data from the seven cases were analysed together (Proportion of correct responses: Amplitude modulated (AM) sound, IM vs NIM 0.99±0.01 vs 0.98±0.01, F_(1,6)_=4.03, p=0.09. Narrow band noise (NBN), IM vs NIM 0.99±0.01 vs 0.96±0.05, F_(1,6)_=1.76, p=0.23. Silence: IM vs NIM 0.92±0.06 vs 0.91±0.08, F_(1,6)_=0.05, p=0.84).

Animals were able to detect AM and NBN stimuli but less able to identify silence (Fig 1E and S1E). Animals with tinnitus might be expected to confuse NBN stimuli with an internally generated percept and therefore show a bias towards responding during silence trials as if NBN stimuli had been presented (Fig S1D).

At both timepoints following NOE, NBN detection performance on this task was similar to baseline (NBN, BL vs Post1, 0.97±0.04 vs 0.97±0.02, β=0.01,t_(16)_=0.32,p=0.76, BL vs Post2, 0.97±0.04 vs 0.91±0.1, β=-0.03, t_(16)_=-2.19, p=0.05) (Fig 1E), whereas a significant decrease in AM detection performance was evident six months later (AM: F_(2,14)_=6.57, p=0.01, BL vs Post1, 0.99±0.01 vs 0.99±0.01, β=-0.001, t_(16)_=-0.19, p=0.86; BL vs Post2, 0.99±0.01 vs 0.98±0.01, β=0.004, t_(16)_=2.39, p=0.03). The animals also achieved lower scores for silence trials following NOE, but this difference was not significant (Silence: BL vs Post1, 0.91±0.07 vs 0.8±0.83, β=-0.17, t_(16)_=-2.15, p=0.05; BL vs Post2, 0.91±0.07 vs 0.83±0.06, β=-0.06, t_(16)_=-0.87, p=0.4) (Fig 1E). However, note the relatively high variability in the NBN and silence trials, also in line with previous studies using the same paradigm [49,55].

Response times for both correct and incorrect trials, although initially unchanged in the trials in which auditory stimuli were presented (AM: BL vs Post1, 1.38±0.12s vs 1.5±0.21s, β=0.03, t_(16)_=2.2,p=0.05; NBN: BL vs Post1, 1.43±0.1s vs 1.49±0.16s, β=0.02, t_(16)_=1.21,p=0.25), became significantly longer six months after NOE (AM: BL vs Post2, 1.38±0.12s vs 1.77±0.26, β=0.09, t_(16)_=6.61,p<0.001; BL vs Post2, 1.43±0.1s vs 1.94±0.47s, β=0.1, t_(16)_=2.23, p=0.04) (Fig S2C and D). In silence trials, longer response times were initially present in ‘correct’ trials (BL vs Post1, 3.16±0.54s vs 3.95±0.95s, β=0.1, t(16)=3.08, p=0.01) (Fig S2C), but had returned to baseline levels six months after NOE (p=0.1) and were unchanged in ‘incorrect’ trials (p>0.1) (Fig S2D). Notably, response times in silence trials six months after NOE were nearly identical between animals, which may indicate a stereotyped response and possibly impaired decision making given an age effect is unlikely (see Methods).

The increased response times after NOE in trials where a sound stimulus was present may, in combination with the small bias of the animals to respond as if a sound was presented in silence trials, indicate increased uncertainty in the animal’s perception about whether a sound was present or not. It seems unlikely that animals had difficulty discriminating between the sound stimuli (AM and NBN) as performance was not affected for NBN trials and only minimally so for AM trials (proportion of correct responses, Fig 1E). Therefore, the long–term effects seen in this paradigm are consistent with the perception of a phantom sound.

To assess the integrity of the auditory brainstem over time, ABRs were measured in all animals one week after NOE (see Methods; Fig 1F). In agreement with previous studies in ferrets [56–58], ABRs presented high variability across individual animals, which, compared to rodents, is likely due to the increased thickness of the skull. However, ABRs showed robust and reliable peaks and troughs with highest sensitivity between 8 and 16 kHz (Fig 1G), corresponding to the highest sensitivity in the ferret audiogram [59].

Following NOE, ABR thresholds significantly increased for stimuli with 8 kHz centre frequencies, the NBN NOE stimulus (BL vs Post1, β =15.71, t_(16)_=7.24, p<0.05), and above (16 kHz, (BL vs Post1, β= 15.71, t_(16)_=2.17, p<0.05) (Fig 1G, Table 1). In the late assessment (Post2), thresholds were significantly elevated across all tested frequencies (p<0.05), except for 1 kHz NBN, suggesting a long–term degradation of auditory function.

**Table 1.**
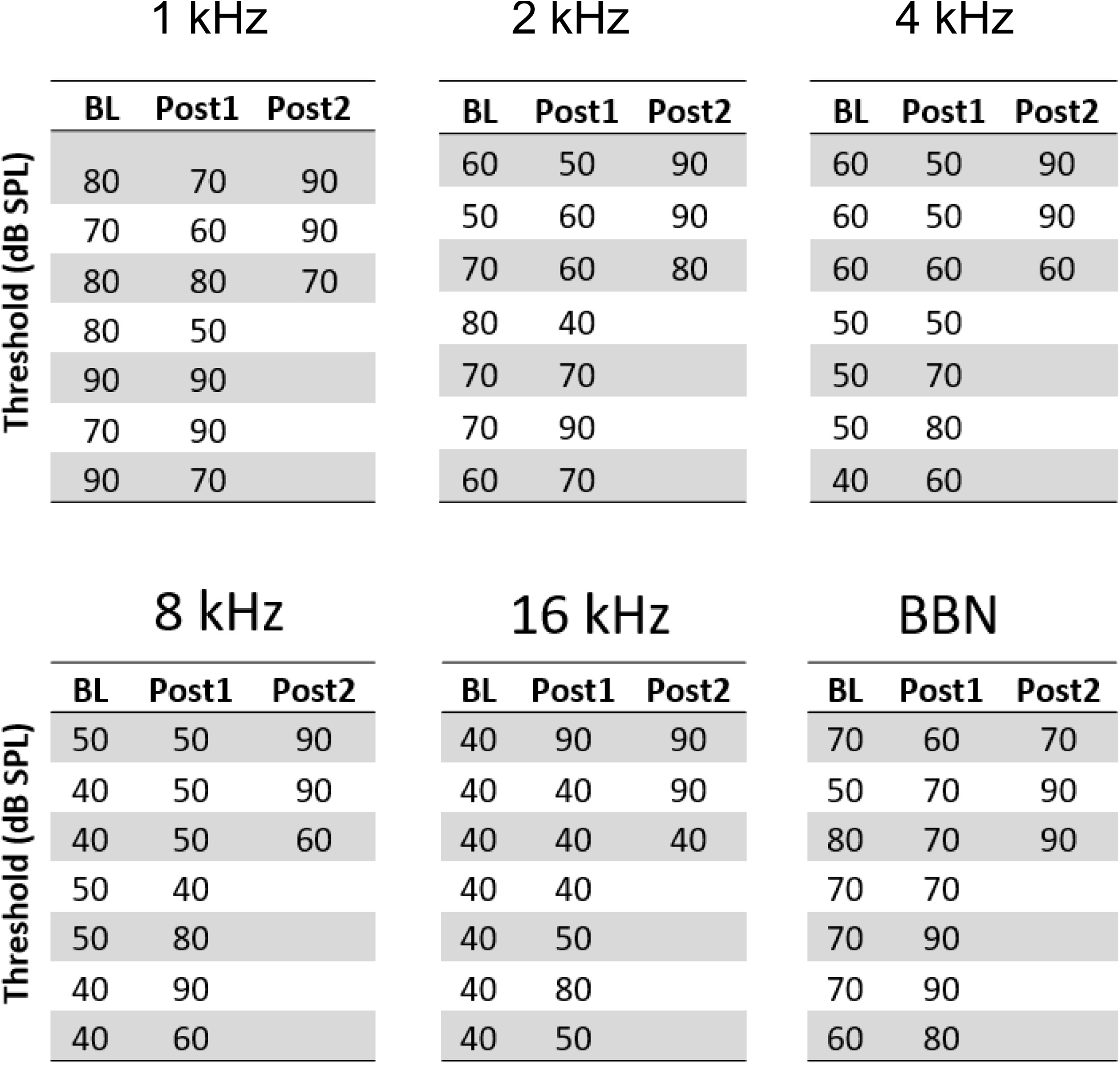
ABR thresholds across time. Different columns, from left to right, correspond to the thresholds of Ferret 1, 2, and 3. Each table shows thresholds for a given stimulus (1, 2, 4, 8, 16 kHz centred NBN and BBN). Each row depicts data from one animal. Note that only in a subgroup of ferrets (n=3), Post2 was assessed.

| 1 kHz |  |  |  | 2 kHz |  |  |  | 4 kHz |  |  |
| --- | --- | --- | --- | --- | --- | --- | --- | --- | --- | --- |
| Threshold (dB SPL) | BL | Post1 | Post2 | BL | Post1 | Post2 |  | BL | Post1 | Post2 |
|  | 80 | 70 | 90 | 60 | 50 | 90 |  | 60 | 50 | 90 |
|  | 70 | 60 | 90 | 50 | 60 | 90 |  | 60 | 50 | 90 |
|  | 80 | 80 | 70 | 70 | 60 | 80 |  | 60 | 60 | 60 |
|  | 80 | 50 |  | 80 | 40 |  |  | 50 | 50 |  |
|  | 90 | 90 |  | 70 | 70 |  |  | 50 | 70 |  |
|  | 70 | 90 |  | 70 | 90 |  |  | 50 | 80 |  |
|  | 90 | 70 |  | 60 | 70 |  |  | 40 | 60 |  |
| 8 kHz |  |  |  | 16 kHz |  |  |  | BBN |  |  |
| Threshold (dB SPL) | BL | Post1 | Post2 | BL | Post1 | Post2 |  | BL | Post1 | Post2 |
|  | 50 | 50 | 90 | 40 | 90 | 90 |  | 70 | 60 | 70 |
|  | 40 | 50 | 90 | 40 | 40 | 90 |  | 50 | 70 | 90 |
|  | 40 | 50 | 60 | 40 | 40 | 40 |  | 80 | 70 | 90 |
|  | 50 | 40 |  | 40 | 40 |  |  | 70 | 70 |  |
|  | 50 | 80 |  | 40 | 50 |  |  | 70 | 90 |  |
|  | 40 | 90 |  | 40 | 80 |  |  | 70 | 90 |  |
|  | 40 | 60 |  | 40 | 50 |  |  | 60 | 80 |  |

The temporally frequency-specific ABR impairment following NOE suggests that behavioural changes affecting wider frequency ranges surrounding the NOE stimulus (such as reduced false alarm rate in operant silent gap detection) cannot be entirely ascribed to hearing loss. Therefore, the animals likely developed an initial NOE–frequency specific hearing impairment and eventually hearing loss for the NOE stimulus and adjacent frequencies, but also tinnitus affecting a wider range of frequencies in auditory silent gap detection.

The seven animals presented specific behavioural and hearing impairment without any noticeable general change in demeanor or wellbeing following noise overexposure. No changes in body weight, bowel habits, fur aspect, or social interaction were observed that could indicate potential noise overexposure related distress (see Methods). Although ABRs were not performed in sham-operated control cases, previous results indicate no differences in auditory cortical responses in adult ferrets of the same age as in the current study, implanted 30 months before auditory brainstem recordings were obtained [44].

### Sleep-wake architecture following noise overexposure

To assess changes in sleep–wake distribution in parallel with the emergence of tinnitus after NOE (Fig 2), three adult female ferrets were implanted chronically with EEG electrodes (frontal and occipital derivation, following standard configuration[36,45,60] (see Methods, Fig 2C), and brain activity was continuously recorded in freely behaving animals for periods of 48 h under baseline conditions, and, as before, one week and six months following NOE (Fig 2A and 2B). We identified four different vigilance states: wakefulness, NREM sleep, REM sleep and a previously described secondary REM sleep, REM2 [45] (Fig 2D and 2E).

**Fig 2.**
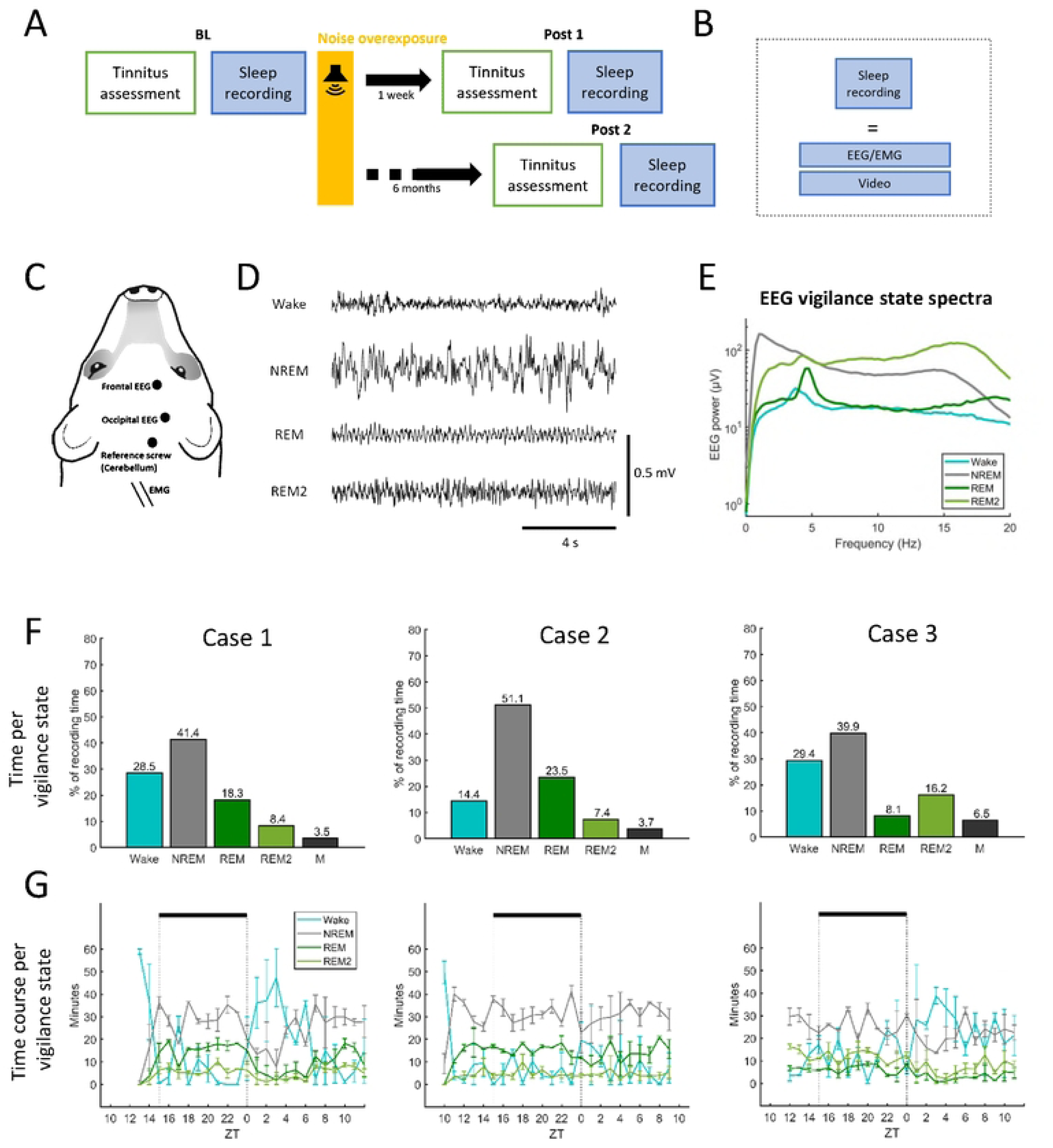
Chronic recordings during sleep and wakefulness. **(A)** Experimental timeline. Ferrets were assessed for tinnitus, hearing loss, EEG brain activity and sleep-wake behaviour before and after noise overexposure (NOE). Assessments of all these metrics took place once under baseline conditions (BL) and on two occasions after NOE, the first assessment (Post1) commencing one week following NOE and the second (Post2) starting six months following NOE. In addition to the ‘tinnitus assessment’, (Fig. 1), EEG using a frontal and occipital configuration, EMG from nuchal muscles along video recording (**B**) were recorded in the freely behaving animals for approximately 48 hours in each condition (BL, Post1, Post2). (**C**) Positions of implanted EEG electrodes for recordings (frontal and occipital), the ground reference electrode implanted over the cerebellum and the EMG wire electrodes on a schematic ferret head. (**D**) Example EEG traces during wakefulness (Wake), non-rapid eye movement sleep (NREM), rapid eye movement sleep (REM) and REM2 sleep (REM2). EEG signals displayed in this panel are band-pass filtered (0.5-30 Hz) and were obtained under baseline conditions (before NOE). (**E**) EEG vigilance state spectra in the ferret (example based on Ferret 3). Data are means across EEG spectra (bin size 0.25 Hz) of the frontal derivation calculated for two consecutives 24 hour recordings. Shading depicts the standard error. Vigilance states (Wake, NREM, REM and REM2) are colour-coded (see inset figure legend). (**F**) Amount of wakefulness and sleep under baseline conditions for each ferret. M, movement artefacts within sleep episodes. (**G**) Time course of wakefulness and sleep under baseline conditions for each ferret in zeitgeber time (ZT) represent the start of light period.

During undisturbed baseline days, the animals spent most of the time sleeping (71.5%, 85.6% and 70.6%, Fig 2F), which is consistent with similar to previously reported sleep amount in ferrets (70.34±1.69%, [45]). Sleep was dominated by non–rapid eye movement (NREM) sleep in all individual animals (Fig 2F). The remaining sleep time was predominantly spent in REM sleep, except in one animal, which spent more time in REM2 than in REM (Fig 2F). In line with previous experiments [45], animals did not manifest strong diurnality and slept both during the light and dark periods (Fig 2G) although the animals’ activity typically increased after light onset for at least 2 hours (2–7 hours across animals; Fig 2G).

Following NOE, all animals developed behavioural signs of tinnitus, as measured by the tinnitus index, TI (see Methods) (Figs 3A and S3). In two animals, tinnitus was most pronounced six months after NOE. In addition, animals developed progressive hearing impairment, which was most pronounced in the animal with the least evidence for tinnitus (Case 3, Fig 3A, S3).

**Fig 3.**
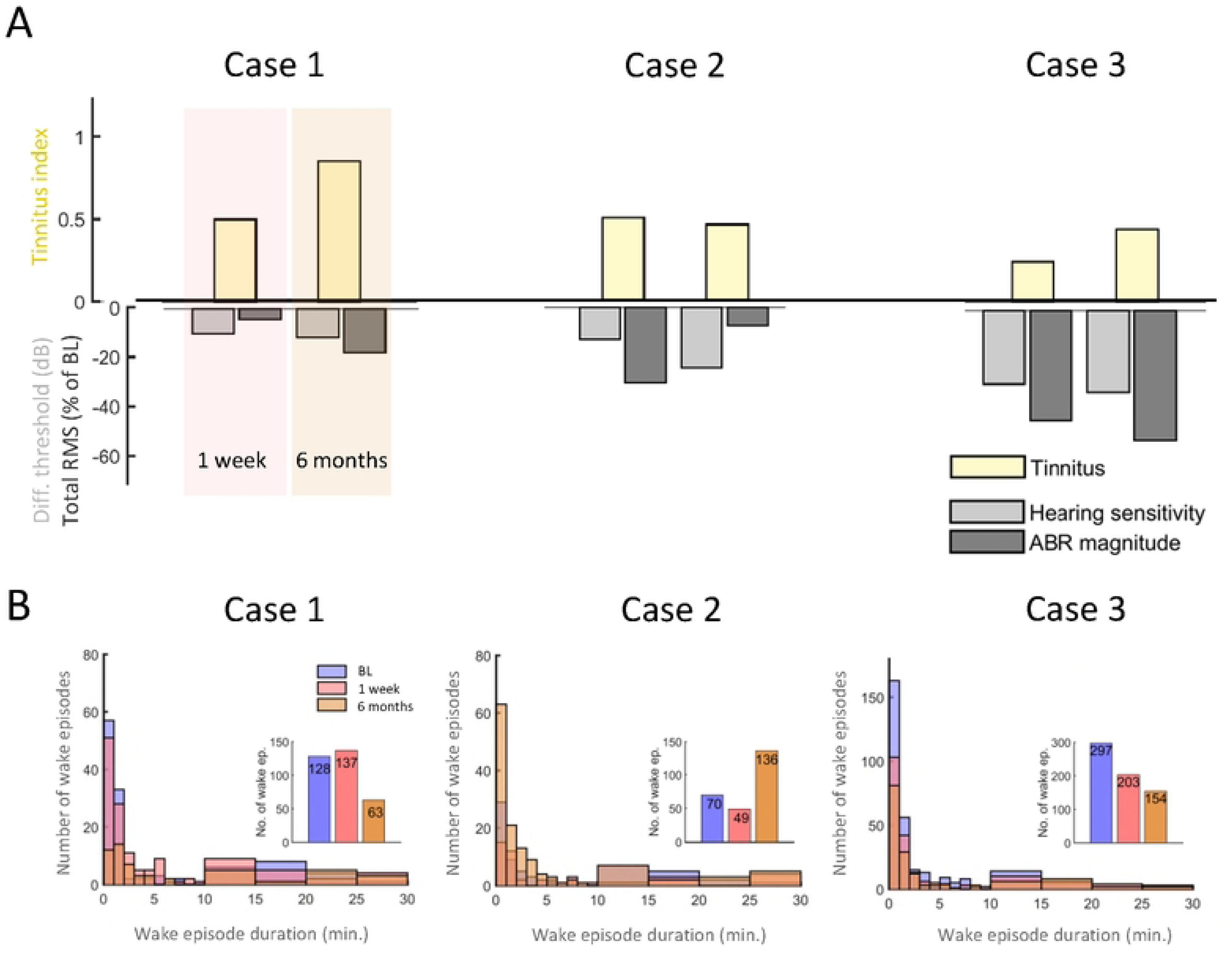
Tinnitus development, hearing loss and sleep disruptions assessed over time. **(A)** Tinnitus and hearing loss development over time in three different ferrets (Cases 1-3). Yellow bars depict behavioural evidence for tinnitus (the tinnitus index, TI, for each animal based on operant gap and silence detection performance; see Methods for details). Grey bars represent the hearing loss after NOE. Light grey bars show the differences from baseline in ABR thresholds while dark grey bars show the change in ABR total RMS magnitude, respectively. The bars are a depiction of the metrics for TI and ABR changes displayed in Fig S3. The size of each bar indicates the change relative to BL of the corresponding measure at Post 1 and Post 2. Each panel represents a different ferret. Light red/light brown shading for Ferret 1 highlight measurements conducted in Post 1 (1 week) and Post 2 (6 months). Plots for Ferrets 2 and 3 have identical layout. Panels left to right: Case 1, Case 2, Case 3. (**B**) Number of wake episodes during 48 hours of baseline recording, one week, and 6 months following noise overexposure. Large panels are histograms depicting the number of wake episodes organised by episode duration and the insets histograms are the total number of wake episodes for baseline (BL), one week (Post 1) and 6 months after NOE (Post 2). The y-axis and the number displayed in the bars in the inset panels depict the number of wake episodes. Note the difference in y-axis scale for Case 3. Note that in Case 2 the number of wake episodes of nearly all durations increases six months following NOE and overlap the other bars (baseline and 1 week post NOE) in the panel.

The sleep pattern changed in all animals following NOE, although this differed between individual ferrets: in both animals with strong signs of tinnitus and weak hearing impairment, sleep became disturbed after NOE but at different time points (Ferret 1 and 2, Fig 3B), while in the animal with weak indication of tinnitus but pronounced hearing impairment (Ferret 3, Fig 3A and 3B) sleep became progressively more stable after NOE, with fewer occurrences of wake episodes during sleep.

To evaluate the individual impact of tinnitus and hearing loss on sleep, each animal was analysed independently with respect to changes in sleep–wake architecture before and after NOE.

### Ferret 1: Progressive tinnitus, mildly raised auditory thresholds and long–term sleep stability

This animal showed marked signs of tinnitus in the first assessment following NOE (TI 0.5, Post1) and a progressive increase towards a TI of 0.9 six months later (Figs 3A and S3). In addition, it showed progressive mild hearing impairment following NOE with threshold elevations of 10 dB (Post1) and 11.7 dB (Post2) (Figs 3A and S3) and a reduction of total ABR magnitude by 4.1% (Post1) and 17.7% (Post2) (Fig S3).

Sleep amount increased transiently following NOE (Figs 3B and S4), in combination with elevated NREM EEG slow–wave activity (SWA, EEG power density between 0.5-4 Hz) (Figs S5 and S6). It is possible that a change in sensory experience following NOE results in compensatory plasticity that is associated with increased sleep need, although increased sleep disruption (Fig 3B) may have contributed to the elevated sleep need in this animal. In the longer term the animal showed a reduction in sleep amount (Wake BL vs Post2, 29.1% vs 34.2% of recording time, Fig S4, grey bars) but also showed reduced sleep disruption (Fig 3B). Note that despite the strong behavioural indication for tinnitus in this animal six months following NOE, sleep amount was largely unchanged as compared to baseline conditions.

### Ferret 2: Stable tinnitus, progressive changes in brainstem activity and long–term disturbed sleep

Ferret 2 showed evidence for stable tinnitus, which emerged soon after NOE (Fig 3A) and was initially (in Post1) of similar intensity to Ferret 1. In parallel, the animal initially showed evidence of increased sleep propensity, with less disrupted sleep than before NOE (70 wake episodes in baseline versus 49 in Post1). However, six months following NOE (Post2), sleep disruption was markedly increased with almost double the number of wake episodes (136 vs 70 wake episodes in BL, Fig 3B), most of which were rather brief (<5 minutes, Fig.3B).

Hearing impairment following NOE was reflected in a progressive ABR threshold elevation (Post1: 13.3 dB, Post2: 25 dB, Figs 3A and S3), suggesting an impairment in auditory sensitivity. Furthermore, total ABR magnitude was temporarily reduced in Post1 (−31.2%), but partially recovered subsequently (Post2: -7.7%, Figs 3A and S3). This later recovery may indicate a long–term compensation following reduced peripheral input through central or peripheral gain elevation.

The initial changes in brainstem evoked activity after NOE were paralleled by a temporary reduction in time spent awake (BL vs Post1: 15.1% vs 12.9%, Fig S4, grey bars) and reduced sleep disruption (Fig 3B). This could be due to temporarily increased sleep pressure following NOE, as also indicated by significantly increased EEG slow–wave activity during NREM sleep (BL vs Post1, Two–way ANOVA, Tukey’s multiple comparisons, p<0.05, Figs S5 and S6) and during REM sleep (BL vs Post1, p<0.05, Fig S5). As in Ferret 1, it is possible that compensatory plasticity after NOE led to increased sleep need reflected in SWA elevation during sleep. The animal’s sleep subsequently became markedly more disrupted, with nearly twice the amount of wake episodes (Post2, Fig 3B) and with overall less sleep than in the baseline condition (Wake BL vs Post2: 15.1% vs 24.0% of recording time, Fig S4). This suggests that the animal may have become more sensitive to external stimuli (hyperacusis) or to internal triggers for arousal, such as tinnitus.

### Ferret 3: Mild tinnitus with progressive, pronounced hearing loss and progressive sleep stability

Ferret 3 showed the most pronounced hearing impairment and the least indications of tinnitus. Even though the TI increased over time, it was generally low (Post1 TI = 0.2, Post2 TI = 0.4, Figs 3A and S3). ABR thresholds were markedly elevated following NOE (Post1: by 30 dB, Post2: by 33.3 dB, Figs 3A and S3) and a progressively reduced total ABR magnitude (Post1: -44.7%, Post2: -52.9%, Fig S3) further indicated the presence of more severe hearing loss in this animal.

There were no signs of increased sleep disruption following NOE (based on the number of wake episodes, Fig 3B). To the contrary, in parallel to progressively impaired hearing, sleep became progressively less disrupted (Fig 3B) and the amount of time the animal spent asleep increased (Wake amount in BL 28.2%; Post 1 20.3%, Post2 23.1%, Fig S4). The decrease in sleep disruption following NOE may be linked to an elevation of the auditory arousal threshold due to hearing loss.

Different from the other animals, sleep in this animal was characterised by lower SWA following NOE than during BL, possibly suggesting more superficial sleep (Figs S5 and S6). The increased sleep amount is unlikely to be a compensatory response to reduced sleep intensity. Instead, this was potentially a consequence of the animal’s ability to maintain consolidated sleep for longer, accumulate a large amount of sleep overall and therefore experience less homeostatic sleep pressure, which is determined predominantly by the time spent awake.

In summary, while all three animals showed a progressive increase in ABR thresholds after NOE, the magnitude of this impairment differed across individuals, as did the emergence of behavioural signs of tinnitus and changes in sleep–wake architecture. Both animals with strong behavioural signs of tinnitus (Ferrets 1 and 2) showed initially higher sleep need after the noise trauma. In the longer term, sleep was disrupted to varying degrees. In the animal with severe hearing impairment and little evidence for tinnitus (Ferret 3), the number of sleep episodes (maintenance) improved following noise overexposure.

### Increased evoked activity in tinnitus is modulated during sleep

To assess whether changes in cortical excitability or responsiveness correlate with tinnitus and might underlie the observed differences in sleep pattern after NOE, auditory evoked activity was evaluated across all vigilance states using free–field sound presentation (see Methods). Briefly, after collecting undisturbed EEG recordings for 24 h, auditory stimuli were presented via a free–field speaker during the subsequent 24 h (Fig 4A). Sounds were one octave narrow band stimuli with centre frequencies of 1, 4, 8 and 16 kHz (820 ms duration, central gap 38 ms, see details in Methods). EEG auditory evoked responses (AERs) were obtained as shown previously in other animal models [61] during wakefulness and in all sleep states under baseline conditions (Figs 4B, C, and S7, S8–10).

**Fig 4.**
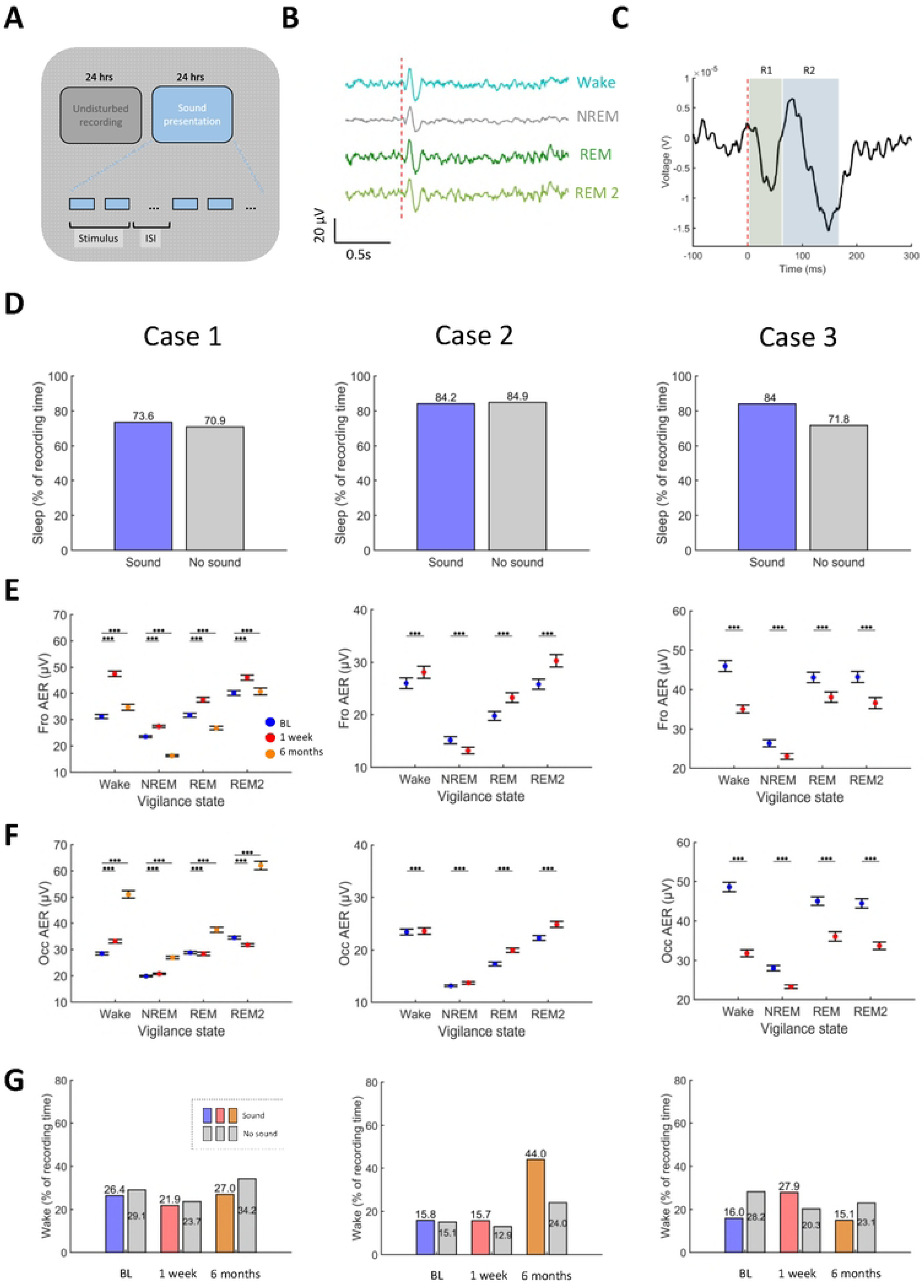
Sound-evoked cortical activity across vigilance states before and after NOE. **(A)** Experimental paradigm. Sounds were presented through a single loudspeaker located at the top of the enclosure over a period of 24 hours subsequent to 24 hours of undisturbed recordings. Sounds were one octave narrow band noise (NBN) bursts centred at 1, 4, 8, and 16 kHz, at a stimulus level of 40, 50, 60, and 65 dB SPL, with a duration of 820 ms that included a silent gap of 38 ms. Stimuli were randomly presented with an interstimulus intervals of 10-42 seconds for a total number of 200 presentations per stimulus-level combination. (**B**) Exemplar average EEG evoked responses (case 3) during Wake, NREM, REM and REM2 sleep during the baseline condition. (**C**) Quantification of EEG evoked response magnitude. Evoked responses of each animal were partitioned into components defined by the negative and positive peaks in the signal. Response magnitude was defined as the difference between a negative peak and the precedent positive peak for each response component (R1 and R2 in this example, see Fig S7 for Cases 1-3). (**D**) Sleep durations (percent of recording time) in undisturbed conditions and with sound presentation measured in basal conditions before NOE. Each panel depicts one ferret. Panels left to right: Case 1, Case 2, Case 3. (**E**) Frontal EEG auditory evoked response (AER) across vigilance states before and after NOE (colour coded). Data are averages across response components, sound level and stimulus type based on bootstrap means ± standard errors (see methods). (**F**) AER for occipital EEG configuration. Asterisks for panels E-F represent statistical significance p values *<0.05, **<0.01, ***<0.001 (GLMM). (**G**) Amount of wakefulness in baseline (BL), Post 1 and Post 2 recordings with sound presentation (coloured bars) and without sound presentation (grey bars), depicted as percent of recording time. In D-G, each panel depicts one ferret; left to right: Case 1, Case 2, Case 3.

All animals spent the majority of time asleep while sounds were presented without marked differences according to whether sounds were presented or not (Fig 4D). These results indicate that the presented sounds were of sufficient intensity to trigger evoked EEG responses but did not disrupt sleep.

There was a statistically significant interaction between the magnitude of the AER (pooled across all response components per animal, Figs 4C, S7, see Methods) and the vigilance state in both the frontal and the occipital EEG (*Frontal EEG*: Ferret1, F_(1,956)_=1.874E+30, p<0.001, Ferret2, F_(1,1915)_=1.364E+31, p<0.001, Ferret3, F_(1,1276)_=1.423E+32, p<0.001; *Occipital EEG* , Ferret1, F_(1,956)_=8.092E+30, p<0.001), Ferret2, F_(1,1916)_=7.802E+24, p<0.001, Ferret3, F_(1,1276)_=4.892E+29, p<0.001). AER magnitudes were lowest during NREM sleep and highest during Wake and REM2 sleep (Figs 4E and 4F, blue symbols), suggesting that NREM sleep may reduce sound–evoked activity. The modulation of AERs by the vigilance state was similar across sound intensities.

When tinnitus was more severe and hearing loss mild (Ferrets 1 and 2, Fig 3A), AERs during wakefulness increased after NOE (Figs 4E, F). This increase in responsiveness was attenuated during sleep, which could explain differences in sleep disturbance (number of wake episodes, Fig 3B) across ferrets with tinnitus. In Ferret 3, which showed pronounced and progressive hearing loss and less severe tinnitus, evoked activity was reduced across all vigilance states after NOE.

### Ferrets 1 and 2: increased evoked activity in tinnitus is reduced during sleep

Ferret 1 showed a progressive increase in evoked activity (pooled across all response components, Figs 4C, S7), mostly in the occipital EEG derivation (Fig 4E,F), alongside behavioural evidence for tinnitus and little change in ABRs (Fig 3A). Elevation of evoked activity was less pronounced or absent during sleep (Figs 4E, 4F and S8).

The initial increase in evoked activity during wakefulness after NOE (Post1) relative to BL was especially pronounced in the frontal EEG derivation (Fig S8) and evident for all stimuli (p<0.001, GLMM). Nevertheless, evoked activity was still significantly elevated in the occipital EEG (p<0.001, GLMM), except for 4kHz NBN (Fig S11, Ferret 1).

The increase in evoked activity after NOE depended on the vigilance state: it was most pronounced during wakefulness in both EEG derivations (Frontal EEG, BL vs Post1, 31.17±0.7 vs 47.53±1.0 µV; Occipital EEG, 28.47+–0.5 vs 33.11±0.7 µV means±SEM; Fig 4E and F). During NREM sleep, the increase was less pronounced (Frontal EEG, BL vs Post1, 23.52±0.35 vs 27.43±0.43 µV; Occipital EEG, 19.88±0.27 vs 20.77±0.26 µV) or, as in the occipital EEG, even absent for REM and REM2 sleep (Fig 4E and 4F). Note that in the frontal EEG, while evoked activity remained lowest during NREM sleep even after NOE, it was still elevated compared to baseline.

In the late assessment, six months after NOE (Post2), the change in frontal evoked activity reversed and AERs for all stimuli approached baseline levels or below in the frontal derivation (Fig S11), which was largely due to drastically reduced AERs during NREM and REM sleep (Frontal EEG: NREM, BL vs Post2, 23.52±0.35 vs 16.3±0.32 µV; REM, BL vs Post2, 31.64±0.74 vs 26.69±0.66 µV, Fig 4E). This could explain why the animal showed prolonged and less disrupted sleep six months after NOE. In the occipital derivation on the other hand, evoked activity increased further for all stimuli (Fig S11). Note that this pronounced increase in occipital evoked activity six months after NOE coincided with the largest behavioural tinnitus index of 0.9 among all animals (Figs 3A and S3) but was not associated with disrupted sleep.

In Ferret 2, AERs were increased relative to baseline in the first assessment after noise overexposure (Post1) (p<0.001, GLMM) at both frontal and occipital EEG derivations (Figs 4E, 4F, and S8) for all sound stimuli (Fig S11). This supports the notion that the gain of auditory evoked responses increased following NOE.

As in Ferret 1, while the increase in evoked activity was evident in both EEG derivations, this was locally modulated across vigilance states: in the frontal EEG signal (measured across all components of the AER), NREM sleep was associated with a reduced evoked response after NOE relative to baseline (BL vs Post1, 15.14±0.7 vs 13.19±0.62 µV, p<0.001, GLMM), whereas in the occipital EEG, the increase in evoked responses was evident in all vigilance states (p<0.001, GLMM, Figs 4E and F). Notably, the reduced frontal NREM AER was present for all frequency stimuli (Fig S11). It is possible, therefore, that NREM sleep had a suppressing effect on frontal evoked activity after NOE.

Six months after NOE, Ferret 2 showed qualitative evidence for further increased auditory evoked activity in both frontal and occipital derivations (Fig S13). Due to decreased signal quality in the Post2 assessment, this could not be quantitatively verified. However, even without sound presentation, the animal showed an ∼60% increase in the amount of time awake in the Post2 assessment (BL vs Post2, 15.1% vs 24.0%, Fig 4G). With sound stimulation, this effect was amplified (time awake, BL vs Post2, 15.8% vs 44%, Fig 4G), while the amount of sleep was reduced (Fig S4), indicating that increased cortical responsiveness may have led to long–term sleep disturbance.

### Ferret 3: generally decreased evoked activity after NOE and profound hearing loss

Ferret 3 showed reduced cortical evoked activity after NOE during all vigilance states (Fig 4E and 4F), in line with marked progressive hearing impairment (Fig 3A).

In the first assessment after NOE, auditory evoked activity (pooled across all response components; Fig S7) was lower than in the baseline assessment (p<0.001, GLMM, Figs 4E and 4F; Figs S10 and S14). This was the case for all sound stimuli and for both the frontal and occipital EEG signals (p<0.001, GLMM, Fig S11). The reduction in AERs was evident during all vigilance states for most stimuli. During REM2 and REM sleep, there were signs of elevated evoked activity for 8 and 16 kHz stimuli, respectively, but only in the frontal EEG signal (Fig S15). Six months after NOE, there were signs of signs of increased evoked activity in the occipital derivation (Fig S14), but this could not be quantitatively analysed due to reduced signal quality.

While the progressive hearing loss in this animal after NOE in undisturbed condition (without sound stimulation) was associated with less time spent awake, a similar trend over time was not apparent with sound stimulation (Fig 4G).

In summary, the two animals that showed greater behavioural signs of tinnitus also developed increased neural evoked activity after NOE although at different time points. The animal with less evidence for tinnitus but severe hearing loss showed reduced auditory evoked activity after NOE. When tinnitus was more severe and frontal evoked activity was elevated, sleep was disturbed, but not when occipital evoked activity was elevated. This might have important implications for the notion of frontal or brain–wide tinnitus representation playing a role in sleep disruption. In both animals with strong evidence for tinnitus, auditory evoked activity was lowest during sleep, suggesting a role for natural brain state dynamics in modulating tinnitus–related activity.

## Discussion

In this case study, we introduced a new animal model of tinnitus, the ferret, which enabled us to track tinnitus development over a period of several months following noise overexposure, concomitant with sleep monitoring. By combining this with the assessment of sleep architecture and spatiotemporal brain activity, this model provided initial evidence supporting the idea that tinnitus emergence, but not hearing impairment, coincides with emergence of sleep disruption. Furthermore, increased auditory evoked activity in tinnitus animals was reduced during sleep, suggesting a potent role for natural brain state dynamics in modulating aberrant brain activity associated with the effects of noise trauma.

In the first part of this study, we characterised the ferret as a novel animal model of tinnitus allowing for the investigation of long–term effects of noise overexposure on behaviour and brain activity. Ferrets offer particular advantages compared to most rodents since their hearing range overlaps with that of humans [62]. Moreover, ferrets can learn sophisticated behavioural tasks [47,63,64], which widens the scope for behavioural tinnitus assessment. The lifespan of ferrets (6–8 years, [65]) and the age of senescence [66] considerably surpass that of mice (1–2 years, [46,67]) and rats (2.5–3.5 years, [68]). This allows for longitudinal assessment without age–dependent degeneration, which is essential for investigating the time course of persistent tinnitus and its comorbidities. Furthermore, as shown in this study, ferrets are readily trained on operant tasks suitable for detecting the presence of tinnitus.

We demonstrated that noise overexposure in ferrets is not only associated with hearing impairment, but also with changes in behavioural performance that are indicative of tinnitus. Further, we show that measures of the degree of tinnitus and hearing impairment following noise overexposure are highly idiosyncratic, in line with variable effects of noise trauma seen in other animal models [69] and the tinnitus heterogeneity characteristic for humans [70]. Our findings show that the ferret could provide a potent model for studying persistent tinnitus on a case–by–case basis and for assessing aspects of tinnitus that have so far been beyond reach due to the limitations of the animal models used and the constraints of human studies. Future work with larger groups with tinnitus, hearing loss alone, and tinnitus plus hearing loss should lead to the consolidation of the ferret as a valuable animal model in tinnitus research.

In the second part of this study, we obtained chronic EEG recordings to investigate changes in evoked activity and sleep–wake pattern in parallel to emerging tinnitus and hearing impairment. Evoked potentials were recorded with EEG electrodes implanted frontally and occipitally, as is standard for sleep recordings [71], which makes it likely that the measured signals originated in the auditory cortex but were possibly influenced by activity in cortico-cortical connections. As in a recent study on tinnitus conducted in macaques [72], we prioritised conducting comprehensive behavioural and electrophysiological paradigms for each animal for an extended period over shorter and less detailed testing of a larger number of animals.

Interestingly, REM2 sleep amount increased in the three cases followed noise overexposure (Post1) regardless of their different initial magnitude of tinnitus, although only transiently for ferrets 1 and 3 (Fig S4). This may suggest that brain activity characteristic for this state, in particular oscillations in the beta and possibly gamma range, may be a marker for consequences of NOE. In humans exhibiting residual inhibition, gamma activity is positively correlated with tinnitus [73]. Future investigation may explore whether changes in cortical activity reflected by the REM2 state in the ferret also reflect initial tinnitus activity during sleep or the initial brain response to the noise overexposure.

We found that the single case developing severe and progressively worse hearing impairment after noise overexposure also developed progressively stable, prolonged and lighter NREM sleep. Building on previous findings showing that cochlear lesions can reduce wakefulness and prolong sleep [74], our results indicate that hearing impairment may increase sleep maintenance and lead to fewer awake episodes, likely as a result of increased sensory disconnection. This differed in cases where tinnitus accompanied hearing impairment. Animals displaying more severe tinnitus following noise overexposure developed reduced and more disrupted sleep. While these results do not demonstrate a causal relationship between tinnitus–related aberrant brain activity and impaired sleep, the parallel emergence of tinnitus and sleep disturbance over time is highly suggestive of such a connection. Indeed, the elevated auditory evoked cortical activity in animals with signs of severe tinnitus supports this possibility.

Although cochlear damage after noise overexposure can lead to a compensatory increase in excitability that restores evoked activity [75,76], previous studies have shown that increased activity and excitability along the auditory pathway may be a correlate of tinnitus [23,77–80]. More specifically, elevated cortical activity has been reported in humans [81,82] and in a range of animal models of induced tinnitus (chinchillas: [80], cats: [83], guinea pigs: [84,85], rats: [86], and mice: [87]).

Even local hotspots of raised cortical activity can have a widespread effect on the sleeping brain due to the extensive interconnectivity between cortical areas [37]. And since stimulation in the auditory modality is a particularly potent trigger for arousal from sleep [88], increased spontaneous and evoked neural activity in tinnitus may not only explain the sleep impairment observed in ferrets with tinnitus in this study, but also the sleep disturbance so widely reported in human tinnitus sufferers [13–21], which may itself contribute to the distress tinnitus sufferers experience. Further support for this notion may be provided by assessing tinnitus–related changes in local brain activity in regions that are sensitive to shifts in vigilance state, such as the auditory cortex [23,78,80]. Manipulations that alleviate tinnitus in animal models, e.g. multisensory [89,90] or vagus nerve stimulation [78,91], before, during or after sleep may uncover the causality in this relationship. The ferret model of noise induced tinnitus will be valuable for detailed long–term investigation and manipulation of tinnitus and will also help to answer the question of whether altered sleep contributes to tinnitus comorbidities, such as distress, depression and anxiety [10].

It is possible that animals displaying signs of tinnitus in our study also developed other noise–induced conditions, such as elevated sensitivity to environmental sounds or hyperacusis [69,87,92]. Although this would not account for all the behavioural deficits observed in this study, such as impaired silent gap detection, it cannot be ruled out that hyperacusis was an additional consequence of noise overexposure and contributes to sleep impairments. Hyperacusis has been suggested as common factor in both tinnitus and insomnia [93].

The results of this study point to a potential role for sleep in the transient relief from tinnitus. Increased evoked activity induced by noise overexposure, which is associated with tinnitus [78], was less pronounced during sleep. Therefore, naturally occurring brain states that are known to interfere with sensory signal processing [3,94] may also mitigate the effects of altered excitability following noise trauma. This may be due to sleep of increased intensity after noise overexposure reflecting elevated sleep drive produced by persistent tinnitus–related brain activation in the waking stage [95–98]. Previous studies indicated that prolonged brain activation raises the internal and network drive of neurons to engage in sleep–specific firing patterns reflected by slow–wave activity [60,99], producing a functional state with the potential to override aberrant brain activity associated with tinnitus [32]. It remains to be seen whether sleep also interferes with aberrant spontaneous activity in individuals with tinnitus.

Following progress in identifying behavioural and physiological changes in awake or anaesthetised animal models of tinnitus [78,79,95,97,100–104], it is now possible to track such objective tinnitus markers, for example via high–resolution recordings in brain regions of interest [78,105]. Extension of such recordings across sleep and wake states should help identify the distinct role of natural brain state dynamics in the modulation of tinnitus markers, especially if sleep is manipulated in parallel, such as through artificial enhancement of slow waves [106,107] or homeostatic increase of sleep pressure after periods of extended wakefulness [30,38,60]. Our results suggest that such approaches for modulating sleep might provide a route towards controlled relief from tinnitus.

Although tinnitus is the most prevalent sensory phantom percept in humans, it is not the only one. The phantom limb syndrome in the somatosensory system (the work of Ambroise Paré reviewed in [6], also [5,7]) and the Charles–Bonnet syndrome in the visual system [108] are well recognised phenomena where phantoms are perceived in the absence of the correspondent sensory stimuli. Since sleep attenuates sensory evoked responses (e.g. [109] in the visual cortex, [3] in the perirhinal cortex), sleep–related modulation of sensory phantoms might extend to multiple modalities.

The most widely accepted theory for the basis of phantom percepts, the deprivation theory [110,111], postulates that a reduction in sensory input is an essential trigger, while more recent findings suggest that further changes in sensory precision and predictive coding are necessary for tinnitus to develop [27]. In both models, the most potent risk factors for tinnitus remain clinically identifiable hearing loss, hidden hearing loss [76,92] and subsequent brain plasticity. Therefore, tinnitus provides a unique model to study brain plasticity outside of the homeostatic range and its relationship with hearing impairments. The dynamics of natural brain states may be a major player in the modulation of either or both conditions and the study of sleep could lead towards new therapeutic avenues in tinnitus, in particular, and in sensory impairments, in general.

## Conclusion

We investigated the interaction between sleep and tinnitus in a novel ferret model of noise overexposure–induced tinnitus. A combination of tinnitus and hearing assessments, vigilance state analysis, and the measurement of spontaneous and auditory–evoked EEG activity across vigilance states provided evidence for a bi–directional interaction between tinnitus and natural brain state dynamics. Cases developing tinnitus also exhibited sleep impairments, suggesting a link between noise–induced tinnitus and sleep disruption. Neural markers of tinnitus were reduced during sleep, suggesting that the sleep state may transiently mitigate tinnitus. This reveals a new angle to tinnitus research, which could prove fruitful in explaining tinnitus comorbidities and offer opportunities for new experimental approaches. Most importantly, these results demonstrate that sleep has the potential to explain and, ultimately, help to mitigate the neural consequences of phantom percepts.

## Declarations

### Availability of data and materials

The raw data and code used in this study are available from https://github.com/l-milinski/Code_for_Milinski_et_al_2023. Scripts and data related to spectral analysis will be made available upon request as the data size is too large for github.

### Authors’ contributions

L.M. conceived, performed and interpreted experiments, analysed the data, and wrote and edited the manuscript; F.R.N. conceived, performed and interpreted experiments, contributed to data analysis and edited the manuscript; M.K.J.E. assisted in performing experiments and contributed to data analysis; A.J.K. contributed to data analysis and edited the manuscript; V.V.V. conceived and interpreted experiments, contributed to data analysis and edited the manuscript. V.M.B. conceived, performed and interpreted experiments, contributed to data analysis, and wrote and edited the manuscript.

## Acknowledgements

We are grateful to Susan Spires and Ana Sánchez for their support in conducting ferret operant behaviour experiments.

